# Thermodynamic projection of the antibody interaction network: the fountain energy landscape of binding

**DOI:** 10.1101/124503

**Authors:** József Prechl

## Abstract

The complexity of the adaptive immune system in humans is comparable to that of the central nervous system in terms of cell numbers, cellular diversity and the network of interactions between components. While the application of molecular biological methods and bioinformatics has brought about an ever deepening and sharpening static description of the molecular and cellular components of the system, a unifying theoretical understanding of the laws governing the dynamics of the system is still lacking.

We have recently developed a quantitative model for the description of antibody homeostasis as defined by the dimensions of antigen concentration, antigen-antibody interaction affinity and antibody concentration. In this paper we develop the concept of a novel thermodynamic representation of multiple molecular interactions in a system, the fountain energy landscape of binding. We show that the hypersurface of the binding fountain corresponds to the antibody-antigen interaction network projected onto an energy landscape defined by conformational entropy and free energy of binding. We demonstrate that thymus independent and thymus dependent antibody responses show distinct patterns of changes in the energy landscape. Overall, the binding fountain energy landscape concept allows a systems biological, thermodynamic perception of the functioning of the clonal humoral immune system.

---

Blood is a massive and critically important extracellular space of multicellular organisms. It is a fluid tissue with cellular, macro- and small molecular components that perfuses the whole multicellular organism, being in direct contact with vascular endothelial cells and blood cells. Its components are potentially derived from any cell of the organism via secretion and leakage (Anderson & Anderson, 2002). Such a hugely diverse molecular pool needs to be regulated with respect to the quality and quantity of its components. One of the mechanisms of regulation is the generation of antibodies by the humoral adaptive immune system (Prechl, 2017a, 2017b). Considering the diversity of antibodies and the diversity of molecular targets, the interaction landscape of humoral immune system is presumably the most diverse in an organism. In this perspective article we approach antibody homeostasis from the thermodynamic point of view, depicting antibody-antigen interactions in a novel energy landscape model. The currently used funnel energy landscape model is suitable for the description of folding and binding of one or a few molecules but it would require landscapes of intractable sizes to depict a whole system, like adaptive immunity. We introduce the fountain energy landscape, a projection of the multidimensional binding landscape of antibodies to the dimensions of entropic penalty and energy of molecular interactions, to accommodate the vast range of interactions of antibodies.

## Energy landscape and antibody binding

Molecular interactions can be described by examining structural, kinetic and thermodynamic properties of the binding. Structural approaches aim to define the relative spatial positions of the constituting atoms of the interacting partners in the bound and unbound forms of a molecule. The advantage of the structural approach is the high resolution visual rendering of molecular structure that helps human perception. Systematic analysis of protein structures gives insight into the evolution of protein complexes and the dynamics of assembly and disassembly (Marsh & Teichmann, 2015). Structural information can also reveal networks of protein interactions (Kiel *et al*, 2008). Kinetic studies follow temporal changes of association and dissociation of interacting partners. These observations are easily applicable to a simple system with a few components only but it is difficult to describe complex systems and crowded molecular environments (Schreiber *et al*, 2009; Zheng & Wang, 2015). Thermodynamics examines the changes in free energy that accompany a binding event; providing statistical descriptions of enthalpic and entropic components of the interaction. Energy landscape theory resolves some shortcomings and integrates these approaches by assuming the presence of many different conformations that converge to thermodynamically stable forms, the route taken to obtain this conformation dictating the kinetics of the events (Bryngelson *et al*, 1995). The intramolecular interactions of proteins lead to the emergence of the functional protein conformation, a process called folding. The energy landscape of folding is assumed to be funnel shaped, the stable form of the protein being at the bottom of the funnel with the lowest free energy state (Wolynes, 2015; Finkelstein *et al*, 2017).

The process of protein folding is obviously strongly dependent not only on general physical parameters, such as temperature and pressure, but on the quality and quantity of molecules present in the system. Water is the solvent of life and interactions with water molecules (Fogarty & Laage, 2014) are of key importance in all molecular interactions associated with life. Additionally, the concentration of hydrogen ions (pH), cations and anions and small molecules in water modulate interactions. Macromolecules influence interactions not only by taking part in the interactions but also by the excluded volume effect, restricting diffusional freedom (Zhou *et al*, 2008). An exact definition of the binding environment in the examined system is therefore indispensable for a realistic depiction of the binding energy landscape. Defining the antibody binding landscape in blood would therefore require at least a complete list of all constituents, and better involve abundance of each molecule.

Antibodies are globular glycoproteins secreted into the blood and other biological fluids by plasma cells (Nutt *et al*, 2015). Antibodies are actually a family of oligomeric proteins, with distinct constant regions that qualify them into classes and subclasses and with distinct variable domains that determine their binding specificity (Schroeder & Cavacini, 2010). While most of us think of antibodies as molecules with a well-defined specificity, in fact the majority of the circulating antibodies (especially of the IgM class) is not monospecific (specific to one target) but rather poly-specific and cross-reactive (Seigneurin *et al*, 1988; Kaveri *et al*, 2012). Any comprehensive systems approach to describe antibody function therefore must account for the presence of both highly specific and poly-specific antibodies. Our quantitative model of antibody homeostasis indeed attempts to provide a unified framework for the whole clonal humoral immune system (Prechl, 2017b). Antibodies are secreted by plasmablasts and plasmacells, descendants of B cells that had been stimulated by antigen. B cells are thus raised in an antigenic environment, the function of the immune system being the selection and propagation of B cells, which can respond to the antigenic environment. The essence of humoral immunity is therefore the definition and control of this antigenic environment by regulating molecular interactions. In thermodynamically terms, the aim of humoral immunity is to generate and maintain a binding energy landscape of antibodies suitable for sustaining molecular integrity of the host organism.

## The binding fountain energy landscape

The funnel energy landscape is a theoretical approach used for the depiction of conformational entropy and free energy levels of one particular molecule (Bryngelson *et al*, 1995). Besides the description of intramolecular binding (folding) it can also be applied for the interpretation of homo- or heterospecific binding, such as aggregation or ligand binding (Zheng & Wang, 2015). If we tried to capture the network of antibody interactions by the binding funnel energy landscape we would face two interconnected problems, one deriving from antibody heterogeneity and the other from target heterogeneity. Antibody variable domains constitute the most diverse repertoire of all the proteins present in the organism, estimates being in the range of 10^9–10^11 different primary structures at any particular time of sampling, the hard upper limit being the number of B cells in a human body, around 10^12 cells (Bianconi *et al*, 2013). Even if the tertiary structures show orders of magnitude of lower diversity we still face an immense variability. On the other side, poly-specific antibodies bind to a multitude of targets, with upper limits to the number of targets being set only by experimentation. A combination of these two factors implies that the binding funnel approach would not allow a clearly comprehensible yet thorough description of antibody-antigen binding. To resolve this issue here we develop the concept of a binding fountain energy landscape model.

Free energy changes associated with molecular interactions can be resolved into components that act against or act in favor of binding. The loss of conformational entropy, a.k.a. entropic penalty, acts against binding while energies of non-covalent bonds (enthalpic component), hydrophobic effect (conformational entropy of water molecules) contribute positively. The net difference between these events determines binding energy and protein stability (Figure 1.). Conformational entropy loss of the antibody molecule thereby sets a minimum energy level that needs to be exceeded for any binding event to be stable. First, let us virtually collect all antibody binding events taking place in our system under examination, blood, and sort these events according to the entropic penalty of binding. For the sake of simplicity let us only consider entropic penalty of the variable domains of the antibodies. Second, let us plot free energy changes against conformational entropy. Since entropic penalty sets a minimum, all stable binding events should appear below the theoretical line representing a gradually increasing entropic penalty (Figure 1.). We can also set arbitrary limits for the free energy decrease, as the range of equilibrium constants for reversible antibody binding are known (Figure 1.B) and we can obtain ΔG from K_eq_ by the equation:

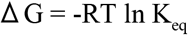

**Figure 1.**
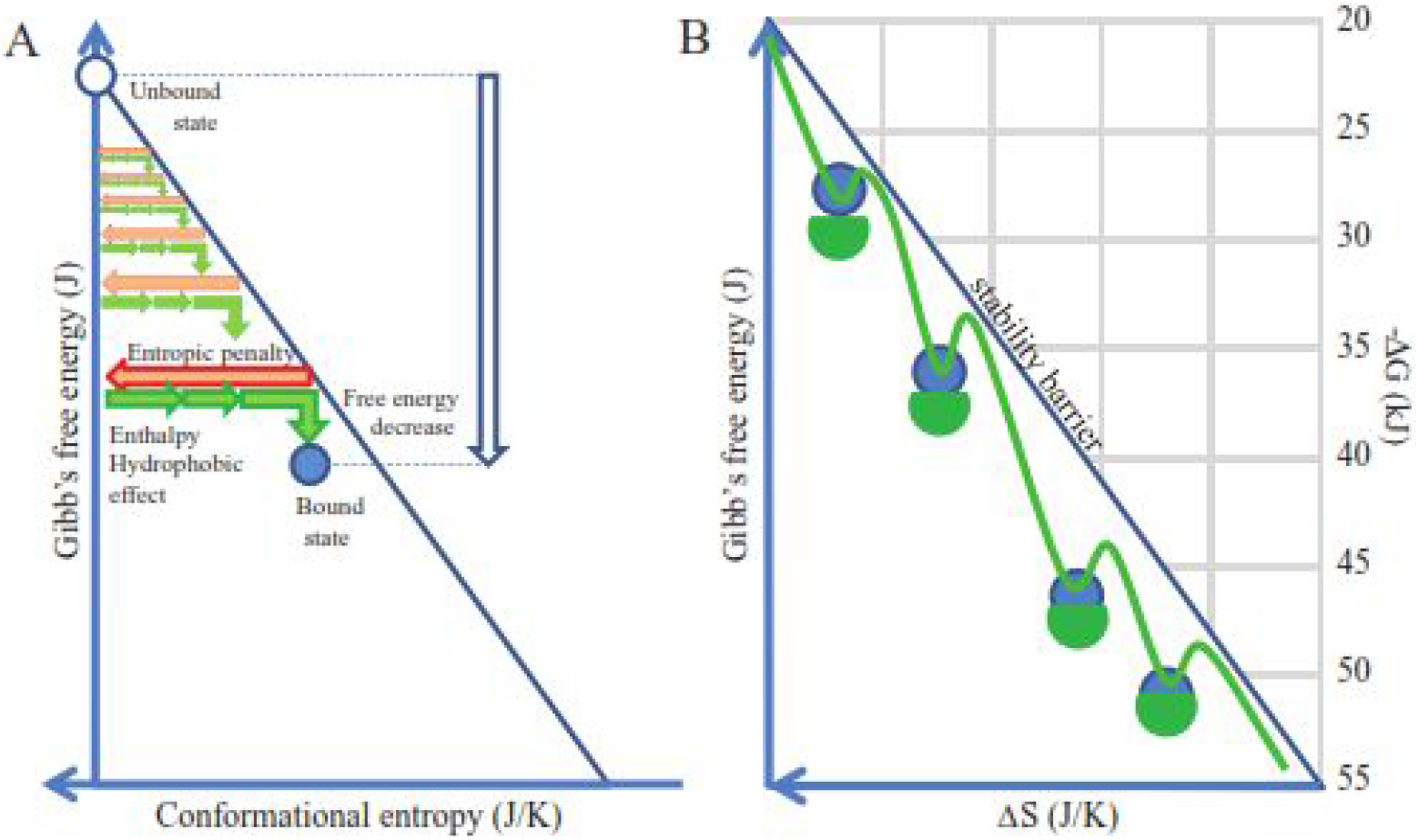
Energy landscape of binding. By anchoring the axis of free energy change at zero entropic penalty we normalize binding events. Any antibody entering the binding landscape appears at the top can search for binding partners with favorable thermodynamic characteristics (A). These binding events take place below the stability barrier, the line representing equality of entropic penalty and energies favoring binding. Theoretically we can collect and position all antibody interactions in this energy landscape (B).

The resulting plot will show the distribution of binding energies against conformational entropy loss. This latter entity is itself associated with the number of atoms at the binding interface and the buried surface area (Marillet *et al*, 2017). Experimental evidence suggests that reversible binding is characterized by a range of energies, limits observed both for maximal and minimal values, which are dependent on the magnitude of the interacting surface, whether characterized by the number of atoms or by buried surface area (Brooijmans *et al*, 2002; Smith *et al*, 2012).

A binding funnel energy landscape focuses on the one native conformation that can be reached via various conformational routes as represented by a hypersurface of conformation, entropy and energy. Instead, we would like to focus on the several different conformational routes taken by several different antibodies while binding to different targets. To this end we assume that all native unbound antibodies enter our landscape at the top of the energy landscape plot. Here conformational entropic penalty is minimal, represents only that associated with folding. Conformational entropy of the antibody, which is the number of different microtrajectories its developing binding surface can take is the highest here. Moving down along any path will lead to loss of free energy and loss of conformational freedom. In order to get a better resolution of the binding landscape let us spin our two-dimensional plot around the energy axis at the maximal entropy to obtain a conical hypersurface in three dimensions (Figure 2). Native unbound antibody molecules entering our landscape will move down along a path while interacting with their targets with an increasing binding energy. This gradual increase in ΔG is accompanied by an increasing involvement of the binding site, called antibody paratope. All stable binding events will take place under the theoretical conical surface generated from the stability barrier line. A binding path ends when the antibody finds its lowest state of energy, corresponding to binding to a target with the highest affinity. Where this point is located depends both on the antibody and the nature of its target (e.g. size, chemical characteristics). The hypersurface of conformations in the space of conformational entropy and free energy generated by this approach we shall call a binding fountain energy landscape.

**Figure 2.**
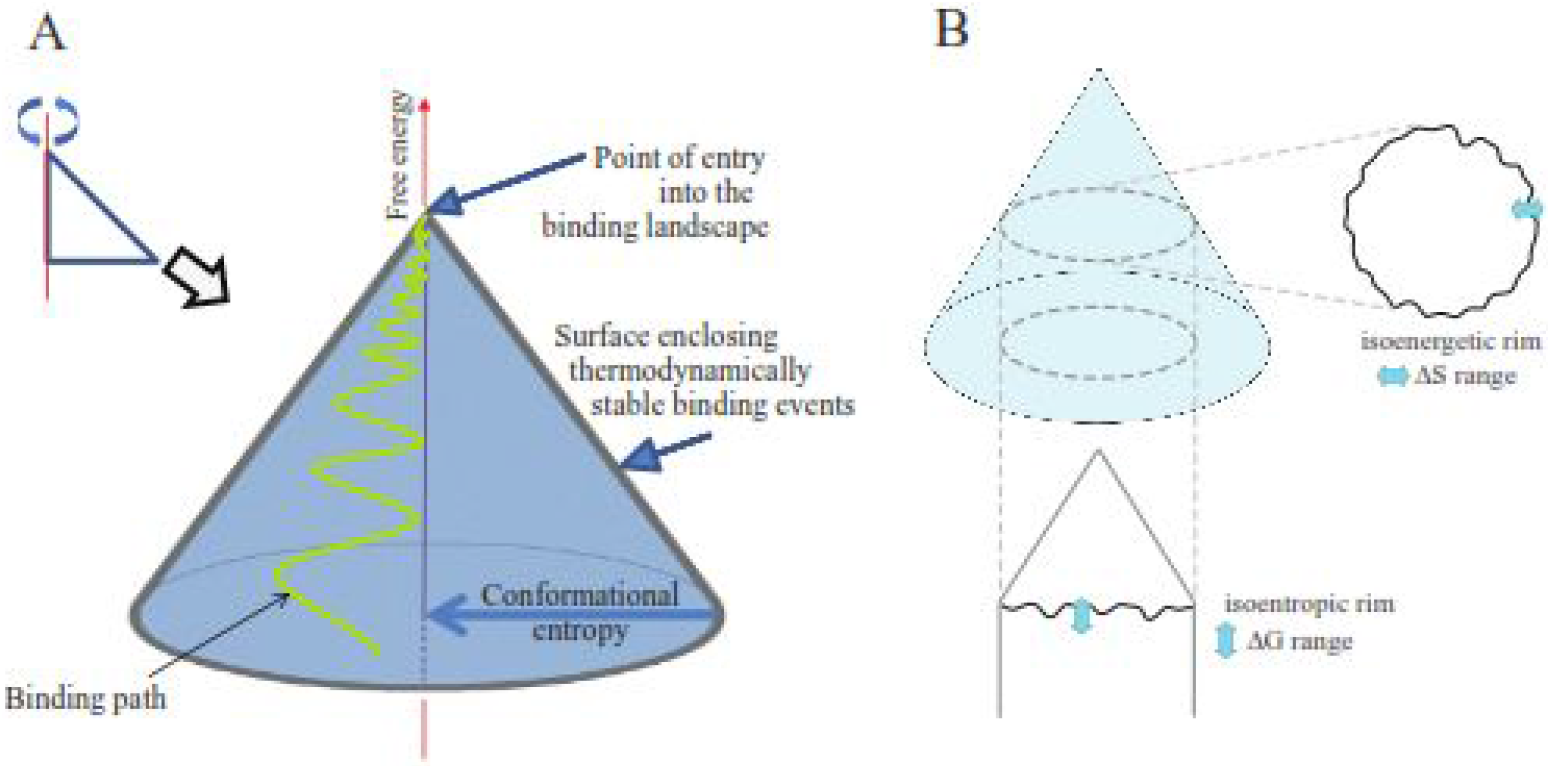
Generation of the binding fountain energy landscape. By spinning the former representation in figure 1 around the anchored axis we obtain a quasi conical surface. This surface encloses all stable binding events of an antibody molecule and is suitable for displaying a binding path (A). Thermodynamically defined subsets of binding events in the binding fountain can be obtained by looking at events in the isoenergetic or isoentropic rim (B).

While the conical surface enclosing stable binding events is a theoretical surface, we can obtain descriptors of real binding events by looking at subsets of events of the interaction space. By cutting the binding fountain horizontally at a given ΔG value we obtain the isoenergetic rim (Figure 2B). The isoenergetic rim is the collection of binding events with identical ΔG and a range of corresponding ΔS. Thus, its ΔS distribution shows the range of entropic penalties that give rise to binding at the given ΔG in our system of study. By cutting the skirt of the cone at a given ΔS value we obtain the isoentropic rim (Figure 2B). The isoentropic rim is the collection of binding events with identical ΔS and a range of corresponding ΔG values. Thus, its ΔG distribution shows the range of free energy changes and corresponding affinity values that give rise to binding at the given ΔS in our system of study. It shows how enthalpy and hydrophobic effects exceed entropic penalty. Please note that as these lines are derived from a hypersurface the lines are theoretical hyperlines themselves, comprising high-dimensional data that cannot be properly visualized in a simple 2D plot.

## Projections of the fountain energy landscape

We have so far worked out an energy landscape interpretation tool, which helps map all the binding events that occur in a molecularly complex environment, such as blood. We assumed that antibodies secreted into the blood gain their native unbound conformations then engage in binding events of various energies until they reach their specific target. The path leading to thermodynamic equilibrium can be rugged, caused by less specific contacts, or smooth, with few intermediate binding states (Figure 3A). It is important to note, however, that blood is the most heterogenous biological fluid, comprising potentially all molecules found in the organism (Anderson & Anderson, 2002). Besides a huge number of secreted molecules any leakage from tissues, debris of cell death and foreign molecules may be present in blood. This vast molecular diversity generates a binding site diversity that we may assume to approach a randomized structural space, representing a huge number of variants of an antibody binding site that covers up to 3000 Å^2^ (Marillet *et al*, 2017). Such a diverse binding space should approach a power law distribution of binding partners, with decay of partners as we increase binding energy or affinity (Figure 3B)(Zheng & Wang, 2015). A rugged start is therefore expectable for all antibodies, with the path smoothing out depending on the paratope properties and the content of the binding landscape. As we approach higher energy and higher entropy loss regions the epitope “sharpens”, as Irun Cohen termed (Cohen & Young, 1991) the gradually increasing affinity of antibodies (Figure 3C). This sharpening involves both a gradually increasing buried surface area and better fitting surfaces and various combinations of these components. It is also apparent that sectors of conformational entropy contain structurally related binding sites, since sharpening reveals more details of epitopes that appear identical at lower resolution (Figure 3D), later maturing into distinct conformational entities. This relationship also reflects the clonal relationship of antibodies going through affinity maturation, gaining sharper but constrained vision of targets by improving their fit (Kang *et al*, 2015).

**Figure 3.**
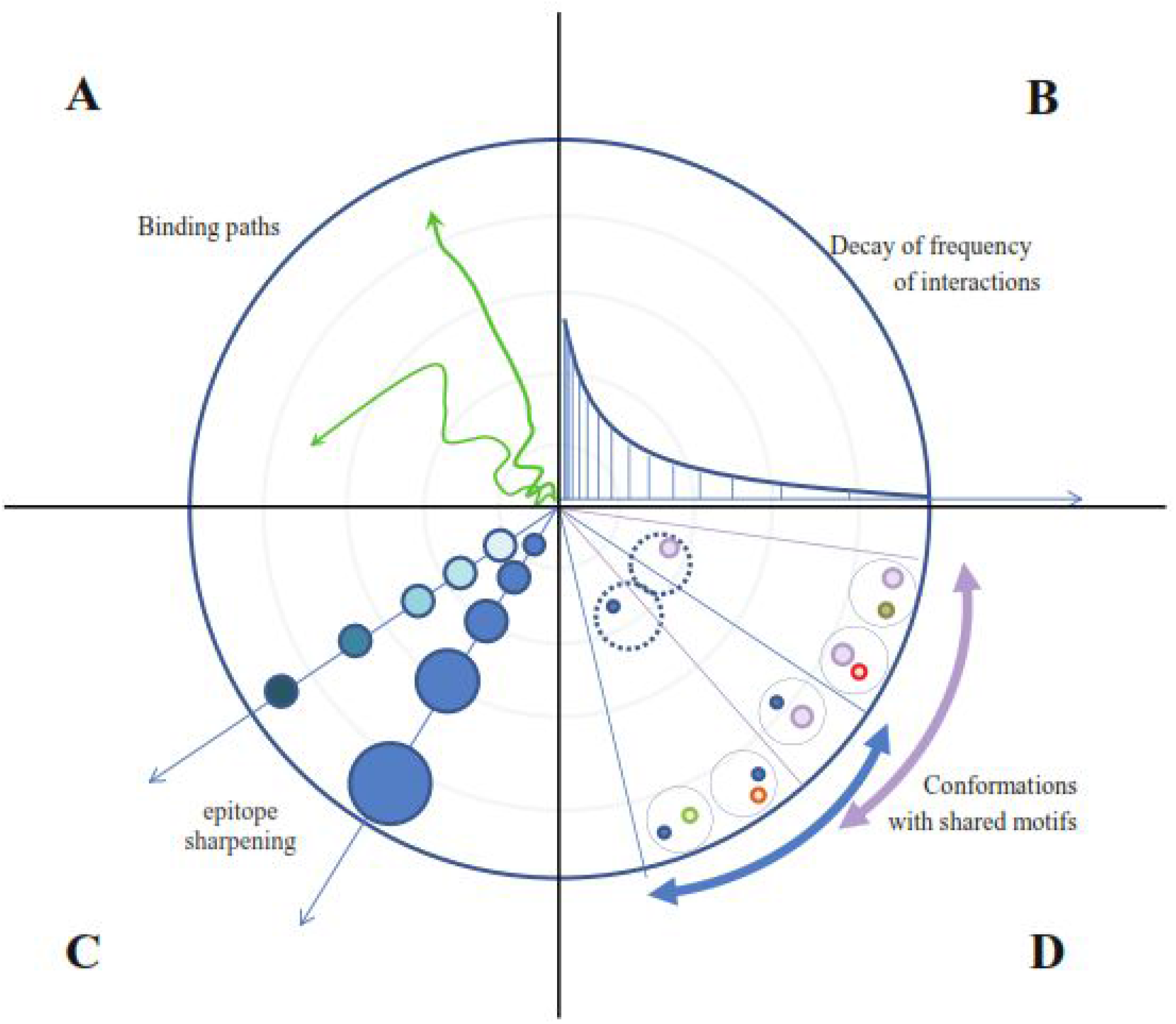
Top view and properties of the binding fountain. Sequential binding of a given antibody appears as a path with more or less rugged track (A). The frequency of interactions decreases by power law decay as we approach high energy binding with high entropic penalty (B). The levels of contact accounting for the entropic penalty increase by improved fit with stronger binding forces and by increased buried interface area (C). Conformations of binding surfaces share common origin with identical structural motifs closer to the “source” of the fountain, the region of low energy interactions (D).

## Interpretation of antibody function as a system of regulated binding landscape

The binding landscape is the set of all potential interactions in a given fluid with given constituents, each interaction being positioned according to the entropic penalty, conformation and free energy decrease. In the binding fountain representation we can trace the fate of a particular antibody in time as a binding path (Figure 4A) or display several different antibodies at an imaginary thermodynamic equilibrium (Figure 4B). Owing to the fact that blood is a highly heterogenous fluid with a vast diversity of potential binding sites the frequency of low energy interactions is very high. At the tip of the fountain antibodies are “surfing” along the ripples of low affinity interactions. Moving down the surface they encounter interaction partners with gradually improved fit, spending more and more time in an interaction, until the target with best fit, that is highest free energy decrease and largest entropic penalty, is found (Figure 4A).

**Figure 4.**
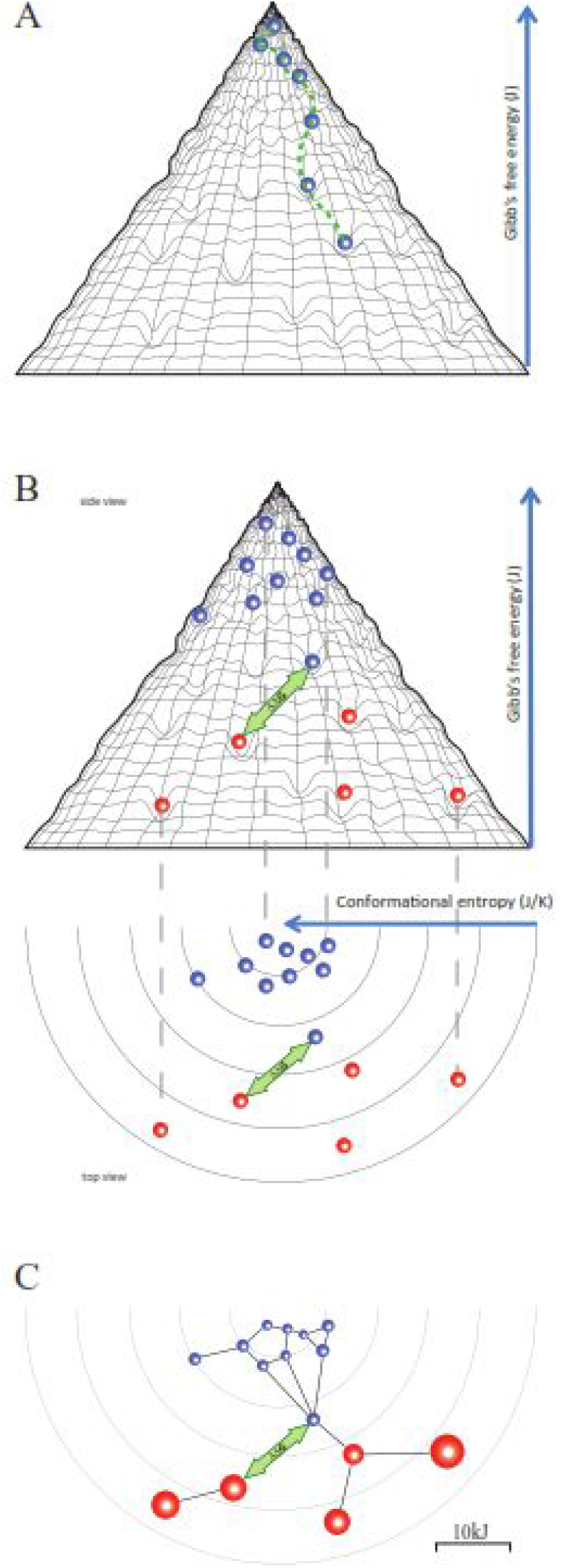
Projections of the binding fountain. An antibody entering the binding landscape engages in serial interactions with increasing energy, taking the molecule down a binding path (A). At an imaginary equilibrium natural antibodies (blue beads) and affinity matured thymus dependent antibodies (red beads) fill the holes of binding, arranged according to their conformation, entropic penalty and free energy level (B). The distance between any two binding events can be expressed as ΔΔG, which represents the cross-reactivity of the two antibodies concerned. We can further project these events into an interaction space where a network is formed based on distance and binding capacity (C).

Interaction in the blood cannot reach thermodynamic equilibrium; molecules are continuously entering and leaving this compartment. On the other hand, because of the constant turbulent mixing the distribution of molecules is constantly approaching homodispersity. Thus, we may display antibodies at an imaginary equilibrium where their position reflects their potential energy minimum in the system. This is where actually target antigen-bound antibody molecules are accumulating (Figure 4B). Registering the position of all the copies of a given antibody species should show a distribution of bound forms determined not only by ΔG but also by the availability of the target molecules, that is antigen concentration [Ag]. The disappearance of the target ([Ag]≈ 0 M) will lead to the disappearance of the low energy position in the landscape. As a consequence, the antibody will accumulate in the interaction with the next available energy level albeit the ratio of bound to free form will be lower as dictated by the higher KD value. Alternatively, the antibody can search the neighboring conformational space along the isoenergetic rim for a binding site with similar ΔG. High concentrations of the target ([Ag]>>KD) will deplete antibody resulting in the potential overflow of related antibodies from the neighboring conformational space. The distance ΔΔG between any two interactions has three components: a free energy component, a conformational component and an entropic penalty component. These components are perceptible from the side view, top view and both views of the binding fountain, respectively (Figure 4B).

As suggested above, besides the presence of targets with a given ΔG the actual concentrations of both Ab and Ag determine the frequency of their interactions and the development of the imaginary equilibrium. To appreciate these factors we can project the interactions of a binding fountain into a space where the distance of the interactions is defined by ΔΔG and the availability of antibody is expressed as the ratio of free antibody to the dissociation constant ([Ab]/KD). This value can be visualized as the radius of the circle representing the interaction (Figure 4C). Please note that this value corresponds to [AbAg]/[Ag], the ratio of bound and free antigen concentrations. This is the network representation of antibody-antigen interactions as we recently described (Prechl, 2017b). A node with a larger radius also implies that the antibody is available for binding to targets in the conformational space in the vicinity of its cognate target molecule. The total combined volume of the nodes of the antibodies under investigation represents the binding capacity of these antibodies in the epitope conformational space.

## The immune response as a regulated binding landscape

The adaptive immune system responds to an antigenic stimulus by the production of antibodies reacting with the eliciting antigen. In our binding landscape antigenic stimulus appears as an impression on the hypersurface representing antibody interactions, the position of the impression being determined by both the conformation of antigen and the conformation of fitting antibodies. The fact that an antigen can stimulate the humoral immune system implies that secreted antibodies that could efficiently bind to the antigen are not present. The antigen therefore binds to the membrane antibodies (B-cell receptors, BCR) of specific B cells (Figure 5). If BCR engagement reaches a threshold the affected B cells proliferate, differentiate and secrete antibodies (Prechl, 2017a). Depending on the nature of the antigen, the route of entry into the host, the presence of costimulatory signals, the ensuing response can proceed basically in two forms. A thymus independent (TI) response will result in the generation of antibodies with binding properties identical to the parental B cell, since there is no affinity maturation. The structure of the binding site does not change, conformation, entropic penalty and ΔG of binding will be identical to the original interaction (Figure 5A). These interactions take place in regions with moderate conformational entropy loss and high interaction frequency, meaning that of the huge repertoire of BCRs several will respond. The response appears as a standing wave, the appearance of antigen showing as the development of the impression, the response of antibody secretion as the disappearance of the impression as free antigen is replaced by bound antigen and immune complexes are removed. This kind of response seems suited for keeping concentrations of target molecules stable. We can think of the response as a closely knit elastic net that regains its original shape after applying pressure to a point (Figure 5A). Thymus dependent (TD) responses will involve the affinity maturation of the antibody binding site, the sequential generation of antibodies with increasing affinity. As the binding site matures the entropic penalty and ΔG increase. The interactions will take place at different positions of the binding landscape (Figure 5B). The response appears as a propagating wave sweeping down the slope of the binding fountain energy landscape. This wave is taking along the antigen, resulting in the efficient elimination of antigenic molecules.

**Figure 5.**
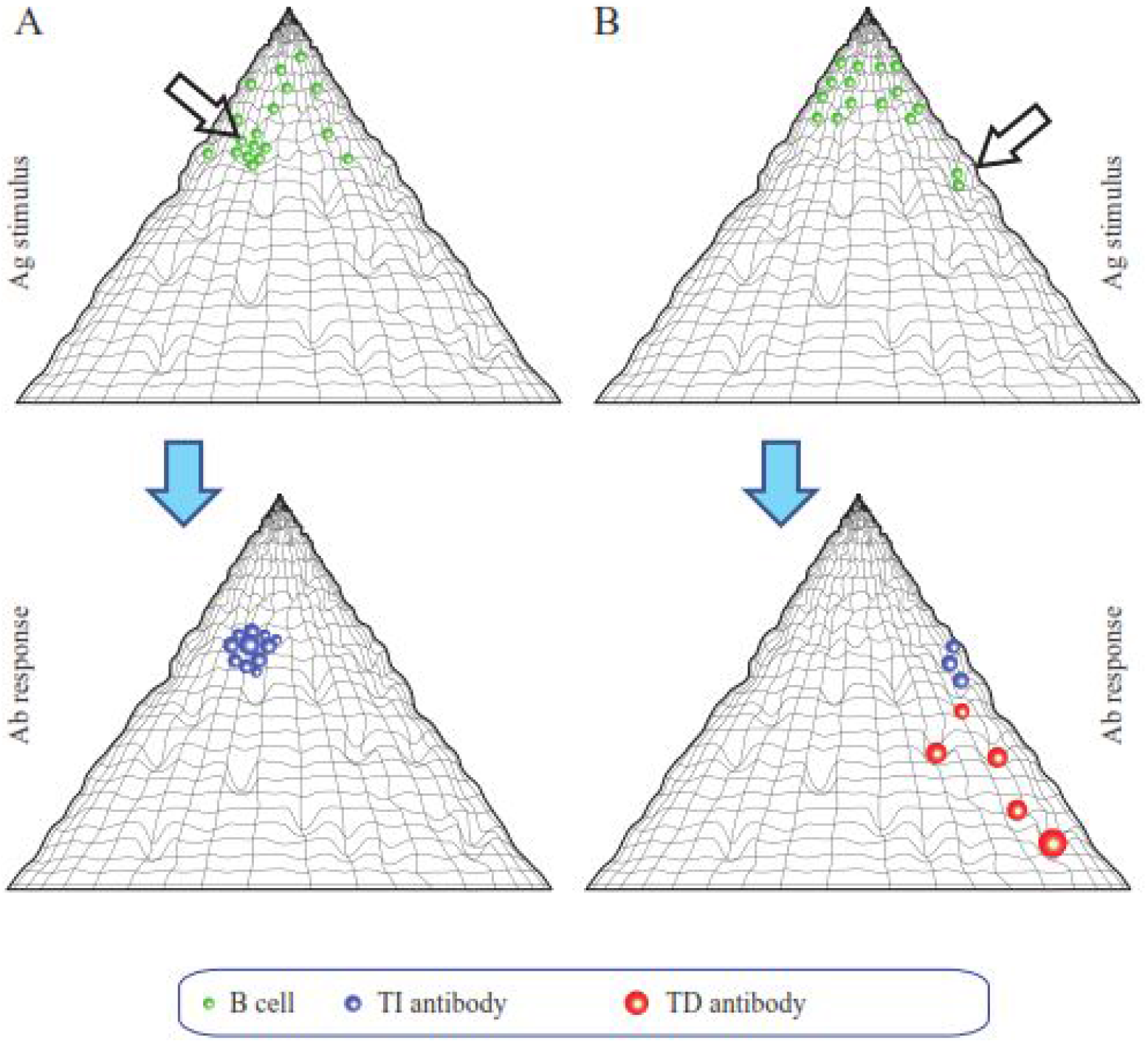
Characterization of fundamental immune response types using the landscape. Thymus-independent responses are characterized by antibodies of lower affinity. A closely knit network of antibody forming cells respond as an elastic net (A). Thymus-dependent responses are characterized by the development of antibodies with increasing affinity. This corresponds to a wave of interactions sweeping down the slope of the fountain (B).

It is important to note the relative identity of binding partners in this landscape: an antibody can bind to antigens but can also be the target of another antibody. The unique binding site of an antibody, the paratope that determines idiotype (identity as a binder), is itself part of the binding landscape. This can be especially important for antibodies with high intrinsic specificity rate (Zheng & Wang, 2015) that are eager to bind and reach their conformation with lowest energy level. We suggest that in the absence of antigen these high affinity binders could be refrained from non-specific binding by engaging their binding sites in lower affinity interactions.

## Proposal

Our current systems biological approach to adaptive immunity can be basically divided into sequence and structure based methods. Next generation sequencing has made it possible to catalogue B-cells as defined by their rearranged heavy and light chain sequences. This approach has very high resolution, allowing the identification of hundreds of thousands of different cells in a sample. The pairing of the chains can also be identified by microfluidic and genetic techniques. Sequencing, in combination with immunogenetic databases and bioinformatic prediction tools, can therefore be used to follow resolve clonality and monitor diversity. The structural and binding properties of reconstituted recombinant antibodies can also be studied albeit on a much smaller scale. One inherent limitation of sequencing, besides intrinsic technological weaknesses, is the sampling bias: we cannot study a whole organism without destroying it, just a sample. In humans this is usually blood. As demonstrated above, the thermodynamic balance of humoral immunity works on a systemic level. Antibodies are produced throughout the body, not just by plasmablast circulating and captured in a blood sample. A systematic study and understanding of humoral adaptive immunity therefore needs different approaches as well.

The second, structural approach is the cataloguing of interactions. Databases devoted to collecting binding data exist but lack quantitative information. The proposed thermodynamic view of the humoral immune system, the network of antibody interactions, suggests that systems level understanding is only achievable by making rigorous quantitative measurements of binding and cross-reactivity. This will require novel technologies and the standardization of antibody and antigen validation. As for technologies, proteins will inevitably follow nucleic acids as subjects of high throughput, high resolution technologies. Standardization of antibody validation is a central issue in life sciences now, and it comes hand-in-hand with antigen standardization. Peptides and mimotopes are the likely candidates for these technologies because of their stability and reproducibility. Exploration of the lower, high-energy binding regions will also require native antigens, since replacement of these would require exact replicas of large interaction surfaces.

The benefits of a systems biological understanding, physical modeling of adaptive immunity would be far reaching. Besides the theoretical beauty observing an evolving cellular system, we expect practical applications in the biomedical fields, from vaccination through autoimmunity and allergy to cancer prevention and treatment.

## Summary

Blood carries potentially all the molecules expressed in the host along with those originating from the environment. To ensure that all these molecules find their intended binding partners a regulated binding landscape evolved: the clonal immune system. The clonal humoral immune system generates a regulated binding landscape by constantly sampling the molecular environment via a huge repertoire of B-cell receptors and by the generation of antibodies with a wide range of specificities and affinities. To allow the thermodynamic representation of this multitude of interactions we show here that this landscape can be visualized as a binding fountain, in an analogy with the folding funnel energy landscape. The binding fountain landscape is an anchored conformation landscape with the conformational entropic penalty of binding anchoring the axis of free energy. Binding sites appear as impressions of a hypersurface, which represents thermodynamically favorable binding events with negative ΔG values. This landscape can be further projected into a multidimensional space of the antibody-antigen interaction network. Systemic perception and interpretation of antibody function is expected to help reveal how the immune system actually functions as a whole, a thermodynamic network of interactions, taking us closer to the understanding of so far underappreciated and less characterized functions of the clonal humoral immune system.

## Acknowledgments

No grant support was used for this study.

## References

Anderson NL & Anderson NG (2002) The human plasma proteome: history, character, and diagnostic prospects. Mol Cell Proteomics 1: 845–867

Bianconi E, Piovesan A, Facchin F, Beraudi A, Casadei R, Frabetti F, Vitale L, Pelleri MC, Tassani S, Piva F, Perez-Amodio S, Strippoli P & Canaider S (2013) An estimation of the number of cells in the human body. Ann Hum Biol 40: 463–471

Brooijmans N, Sharp KA & Kuntz ID (2002) Stability of macromolecular complexes. Proteins 48: 645–653

Bryngelson JD, Onuchic JN, Socci ND & Wolynes PG (1995) Funnels, pathways, and the energy landscape of protein folding: a synthesis. Proteins 21: 167–195

Cohen IR & Young DB (1991) Autoimmunity, microbial immunity and the immunological homunculus. Immunol Today 12: 105–110

Finkelstein AV, Badretdin AJ, Galzitskaya OV, Ivankov DN, Bogatyreva NS & Garbuzynskiy SO (2017) There and back again: Two views on the protein folding puzzle. Phys Life Rev

Fogarty AC & Laage D (2014) Water dynamics in protein hydration shells: the molecular origins of the dynamical perturbation. J Phys Chem B 118: 7715–7729

Kang M, Eisen TJ, Eisen EA, Chakraborty AK & Eisen HN (2015) Affinity Inequality among Serum Antibodies That Originate in Lymphoid Germinal Centers. PLoS ONE 10: e0139222

Kaveri SV, Silverman GJ & Bayry J (2012) Natural IgM in immune equilibrium and harnessing their therapeutic potential. J Immunol 188: 939–945

Kiel C, Beltrao P & Serrano L (2008) Analyzing protein interaction networks using structural information. Annu Rev Biochem 77: 415–441

Marillet S, Lefranc M-P, Boudinot P & Cazals F (2017) Novel Structural Parameters of Ig-Ag Complexes Yield a Quantitative Description of Interaction Specificity and Binding Affinity. Front Immunol 8: 34

Marsh JA & Teichmann SA (2015) Structure, dynamics, assembly, and evolution of protein complexes. Annu Rev Biochem 84: 551–575

Nutt SL, Hodgkin PD, Tarlinton DM & Corcoran LM (2015) The generation of antibody-secreting plasma cells. Nat Rev Immunol 15: 160–171

Prechl J (2017a) A generalized quantitative antibody homeostasis model: regulation of B-cell development by BCR saturation and novel insights into bone marrow function. Clinical & translational immunology 6: e130

Prechl J (2017b) A generalized quantitative antibody homeostasis model: antigen saturation, natural antibodies and a quantitative antibody network. Clinical & translational immunology 6: e131

Schreiber G, Haran G & Zhou H-X (2009) Fundamental aspects of protein-protein association kinetics. Chem Rev 109: 839–860

Schroeder HW & Cavacini L (2010) Structure and function of immunoglobulins. J Allergy Clin Immunol 125: S41–52

Seigneurin JM, Guilbert B, Bourgeat MJ & Avrameas S (1988) Polyspecific natural antibodies and autoantibodies secreted by human lymphocytes immortalized with Epstein-Barr virus. Blood 71: 581–585

Smith RD, Engdahl AL, Dunbar JB & Carlson HA (2012) Biophysical limits of protein-ligand binding. J Chem Inf Model 52: 2098–2106

Wolynes PG (2015) Evolution, energy landscapes and the paradoxes of protein folding. Biochimie 119: 218–230

Zheng X & Wang J (2015) The universal statistical distributions of the affinity, equilibrium constants, kinetics and specificity in biomolecular recognition. PLoS Comput Biol 11: e1004212

Zhou H-X, Rivas G & Minton AP (2008) Macromolecular crowding and confinement: biochemical, biophysical, and potential physiological consequences. Annu Rev Biophys 37: 375–397

